# The correspondence problem: which brain maps are significantly similar?

**DOI:** 10.1101/203083

**Authors:** Aaron Alexander-Bloch, Simon N. Vandekar, Russell T. Shinohara, Siyuan Liu, Theodore D. Satterthwaite, David C. Glahn, Armin Raznahan

## Abstract

A critical issue in many neuroimaging studies is the comparison between brain maps. How should we test the hypothesis that two or more brain maps are partially convergent or overlap to a significant extent? This “correspondence problem” affects, for example, the interpretation of comparisons between task-based patterns of functional activation, resting-state networks or modules, and neuroanatomical landmarks. In published work, this problem has been addressed with remarkable variability in terms of methodological approaches and statistical rigor. In this paper, we address the correspondence problem using a spatial permutation framework to generate null models of overlap, by applying random rotations to spherical representations of the cortical surface. We use this approach to derive clusters of cognitive functions that are significantly similar in terms of their functional neuroatomical substrates. In addition, using publicly available data, we formally demonstrate the correspondence between maps of task-based functional activity, resting-state fMRI networks and gyral-based anatomical landmarks. We provide open-access code to implement the methods presented for two commonly-used tools for surface based cortical analysis. This spatial permutation approach constitutes a useful advance over widely-used methods for the comparison of cortical maps, and thereby opens up new possibilities for the integration of diverse neuroimaging data.

## INTRODUCTION

The spatial dependence in maps of brain activity, morphology, or connectivity impacts statistical inference and the interpretation of neuroimaging studies. It is well-understood that incorrect estimation of spatial smoothing and related statistical tests may result in inflated false negative or false positive rates in functional magnetic resonance imaging (fMRI) analyses (Eklund et al. 2016; Slotnick 2017; Mueller et al. 2017). However, a related issue that has received less attention arises in the comparison between brain maps, i.e., evaluating the possibility that two or more brain maps are partially convergent or overlapping (“the correspondence problem”).

The extent of overlap or convergence between brain maps is a critical issue in many published and ongoing studies of cortical organization. For example, maps of intrinsic resting state fMRI connectivity may overlap with maps of white matter connectivity derived from diffusion imaging data (Honey et al. 2010; Honey et al. 2009; Hagmann et al. 2008; Skudlarski et al. 2008; Horn et al. 2014), or task-based fMRI activation across studies (Smith et al. 2009). Structural covariance, derived from correlations between regions in morphological properties, has largely been interpreted based on the extent of overlap with patterns of intrinsic fMRI connectivity (Kelly et al. 2012; Seeley et al. 2009), white matter connectivity (Gong et al. 2012) and longitudinal maturational coupling (Alexander-Bloch et al. 2013; Raznahan, Lerch, et al. 2011). Shared neurobiological substrates for cognitive functions are often inferred on the basis of overlap between patterns of fMRI activation between different tasks (Otto et al. 2014; Wesley & Bickel 2014; van Belle et al. 2014; Xu et al. 2013), as are shared (or distinct) cellular or developmental origins on the basis of overlap between morphological phenotypes such as cortical thickness and surface area (Maingault et al. 2016; Raznahan, Shaw, et al. 2011). Finally, differences between distinct demographic or clinical cohorts are often inferred in reference to divergent patterns of brain morphology (Douaud et al. 2014) or function (Goksan et al. 2015; Baliki et al. 2014; Zaki et al. 2016). Yet, despite numerous examples where investigators compare spatial patterns between experiments or conditions, there is currently no standardized statistical method for testing convergence.

The manner in which hypotheses about convergence and overlap are tested varies remarkably across the above studies, in terms of methodological approach and statistical rigor. For example, it is not uncommon to simply visualize two maps side by side as evidence of convergence or divergence. Areas of overlap are sometimes highlighted (for example, the conjunction of two independently statistically significant maps). While suggestive, these approaches neglect the possibility, especially for spatially diffuse maps, that such overlap is due to chance and not statistically significant. Statistics such as the spatial correlation between maps are also used to quantify the extent of convergence between maps, and the reported statistical significance of these correlation is in some cases greatly inflated by failing to take into consideration the spatial non-independence of brain maps. In other cases a correction for the spatial degrees of freedom is derived via Gaussian random field theory (Smith et al. 2009; Casanova et al. 2007; Bäuml et al. 2015). Another approach is to use the partial correlation between brain maps after regressing out the shared relationship with anatomical distance (Honey et al. 2009; Horn et al. 2014).

A particular area greatly affected by the correspondence problem is the comparison of community structures (for example, clustering solutions, functional modules, anatomical parcellations or network partitions). The application of community detection algorithms to different datasets is often followed by an attempt to assess whether the resulting community structures are similar or distinct from each other. Examples from the literature include the following comparisons: resting-state and task-based fMRI networks (Kelly et al. 2012); ICA components across different cognitive tasks (Xu et al. 2013); and resting-state modules between clinical groups (Glerean et al. 2016; Achard et al. 2012). Group-wise permutation approaches can explicitly test hypotheses about differences between groups in community structure (Alexander-Bloch et al. 2012), but this approach cannot be extended to the straightforward comparison of the spatial properties of two such structures. Although statistics such as the adjusted Rand index (Hubert & Arabie 1985) and mutual information (Meilă 2007) can quantify the extent of overlap, there is no standard way to interpret the significance of these statistics, which are also influenced by the spatial dependence of brain maps.

A standardized approach to the correspondence problem would help to better address several open questions in neuroscience which hinge on the degree of spatial alignment between different maps of cortical organization. For example, although superficially similar tasks tend to activate similar brain regions (Smith et al. 2009), it remains unclear if and how the brain’s diverse cognitive and affective capabilities are grouped with reference to statistically-significant spatial overlaps in their activation maps. Being able to cluster cognitive tasks into sets that show an overlapping brain activation above and beyond the intrinsic spatial dependence in fMRI data would represent a major step forward in the definition of core functional modules within the brain. A closely related question involves the degree of spatial correspondence between spontaneous activity fluctuations within resting brain (Beckmann et al. 2005) and the coordinated changes in cortical activity that are induced by tasks (Toro et al. 2008). Again, a solution to the correspondence problem would help to resolve this question in a way that controls for the inherent spatial smoothness of shifting brain activity during both rest and tasks (Smith et al. 2009). Furthermore, a rigorous means of quantifying how statistically surprising any given correspondence is, would make it possible to distinguish between brain networks that show differing degrees of alignment between rest and task-induced states. These issues are central to the detection and definition of a core set of dissociable brain networks for targeted investigation on clinical and basic neuroscience (Fornito & E. T. Bullmore 2015). Finally, standardized techniques to control for spatial smoothness in the comparison of brain maps would directly inform longstanding questions regarding the correspondence (or lack thereof) of functional and macroanatomical boundaries within the cortex. To date, this correspondence has been examined for a selected sub-set of cognitive tasks and macroanatomical gyral boundaries (Frost & Goebel 2012), and would benefit from examination throughout the entire cortical sheet across a wide range of cognitive domains simultaneously.

In this paper, we address the correspondence problem using a spatial permutation framework to generate null models of overlap, by applying random rotations to spherical representations of the cortical surface. Our work builds upon initial implementations of spatial permutation (Vandekar et al. 2015; Gordon et al. 2016), in three notable directions. First, we demonstrate the applicability of spatial permutation methods across multiple varieties of correspondence tests, including (i) the relationships amongst meta-analytic patterns of functional activation as defined by the Neurosynth platform (Yarkoni et al. 2011), and (ii) the relationship between canonical cortical parcellations defined with reference to resting state functional connectivity (Yeo et al. 2011) vs. gyral based anatomy (Desikan et al. 2006). The former of these analyses provides a means of determining if sub-groups of cognitive functions (Poldrack & Yarkoni 2016; Poldrack et al. 2011) are clustered in terms of their anatomical substrates above and beyond potential confounding effects of spatial dependence - thereby providing a powerful synthesis of structure-function relationships as charted across decades of functional neuroimaging literature. The latter of these analyses addresses a long-running debate regarding the inter-relationship between morphological and functional boundaries in the human cortex (Ronan & Fletcher 2014). Collectively, the Yeo atlas and Desikan atlas have been used in over four thousand prior neuroimaging studies (PubMed, 2017), making it especially valuable to understand their relationship with each other and with spatial patterns of cortical activation across diverse cognitive tasks (Yarkoni et al. 2011).

Finally, we present a method to test the significance of the overlap between two community structures, which meets a growing need given the rapid proliferation of network-based approaches in neuroimaging science. The ability to ask if and how any two parcellations or modular depictions of the brain are aligned would more effectively exploit the current diversification of available imaging modalities and techniques for parcellation, clustering and community detection (Eickhoff et al. 2017). The code to perform such analyses has been made accessible within two popular pipelines for surface-based cortical analysis (FreeSurfer and CIVET, URL upon publication), so these methods can be easily applied and extended in future work.

## METHODS

### Data

Meta-analytic patterns of functional activation were derived using Neurosynth (http://neurosynth.org) (February, 2015 release), which includes automated meta-analyses of imaging coordinates associated with >3,300 terms in >10,900 studies. We focused the analysis on cognitively relevant maps by filtering for terms that are included in the cognitive atlas (http://www.cognitiveatlas.org/concepts/) (Poldrack & Yarkoni 2016; Poldrack et al. 2011), which resulted in 120 terms. We used the reverse inference maps, comprised of z-scores corresponding to the likelihood that a term is used in a study given the presence of activation in a region (Yarkoni et al. 2011). Compared to the forward inference maps (corresponding to the likelihood that a region reported is active in studies that include a given term), the reverse inference maps are more selective in excluding regions that are diffusely involved in most cognitive tasks. These maps were each projected onto the FreeSurfer average surface by nearest neighbors interpolation (fsaverage5, which contains 10242 vertices per hemisphere), using a mid-gray surface, 50% of the way in between the white and pial surfaces.

As benchmark representations of anatomical regions and functional networks, we used the Desikan Atlas (Desikan et al. 2006) and the Yeo Atlas (Yeo et al. 2011), respectively. The Desikan Atlas, distributed with FreeSurfer (v5.3.0), was derived by manually identifying 34 cortical regions of interest using gyral-based landmarks in 40 human brain MRI scans. The Yeo Atlas was derived from a mixture model of 1000 resting-state fMRI scans after scans were aligned using surface-based realignment. Each vertex was assigned one of 7 resting-state networks.

### Rotational permutation

Spatial permutation of brain maps was performed using angular permutations of spherical projections of the cortical surface. The coordinates corresponding to all of the vertices were rotated at angles uniformly chosen between zero and 360 degrees, about each of the x (left-right), y (anterior-posterior) and z (superior-inferior) axes. For statistical comparisons that necessitated a direct correspondence between the permuted vertices and the original vertices, the original coordinate space was used by assigning the value of the nearest neighbor (based on Euclidean distance) of the vertices in the rotated coordinate space. For opposite hemispheres, the same rotations were applied about the left-right axis of rotation, while opposite (negative) rotations were applied about the superior-inferior axis and the anterior-posterior axis, in order to preserve the contralateral symmetry of the rotated maps. When non-cortical regions (mid-cut or corpus collosum) were rotated into cortical space, these vertices were not included in the calculation of the test statistic (the “correspondence statistic”). We use several different correspondence statistics depending on the specific experimental context, however, the significance of these statistics are always derived relative to the empirical distribution determined by the spatial permutation procedure. Please see Figure 1 for a schematic of the permutation method, as applied to the Desikan Atlas.

**Figure 1.**
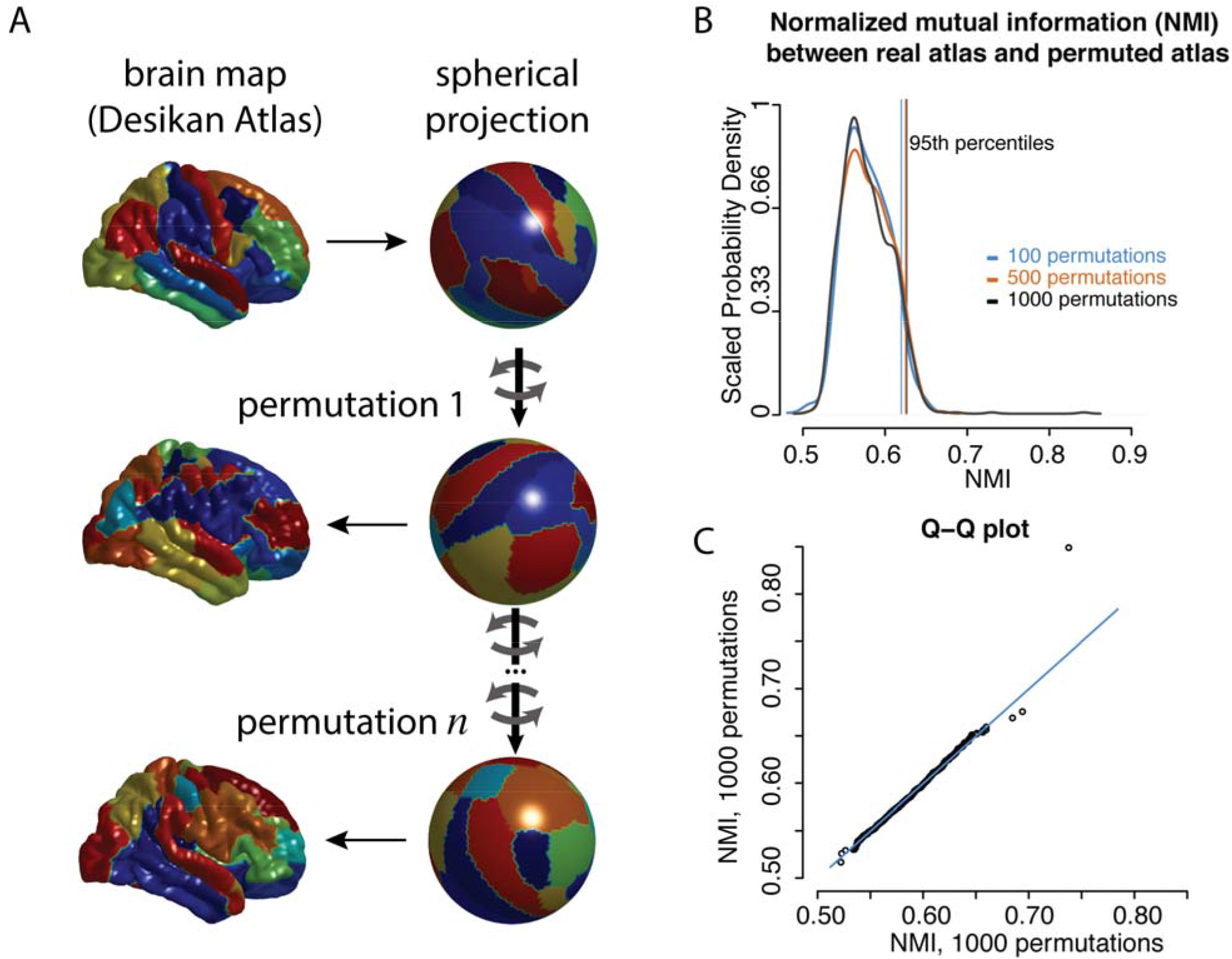
A schematic of the permutation procedure. A) As an illustration, the Desikan atlas is shown in the original space (top left) and spherical space (top right). Each color corresponds to different regions. The spherical coordinates are rotated (mid right, bottom right) and the projected back onto the anatomical surface (mid left, bottom left). B) The degree of similarity between the original parcellation and the rotated parcellations were estimated using the normalized mutual information (NMI). The probability density distributions of this statistic are shown for 100, 500, and 1000 rotations, as well as lines marking the 95^th^ percentile of each distribution. C) A Q-Q plot of the two independent distributions of 1000 rotations each.

### Cognitive clusters of meta-analytic patterns of activation

To quantify the degree of relatedness between the patterns of activation for the 120 cognitive terms, the correspondence statistic was Pearson’s r between every map and every other map, as represented by a 120×120 correlation matrix. These correlations were visualized using complete linkage hierarchical clustering, with the distance between the maps calculated as 1 − r. To test the statistical significance of these correlations, we generated 1000 rotational permutations of the data as described above. For each permutation, we generated a 120×120 permuted correlation matrix (the correlations between the permuted data and the original data). The maximum of the absolute values of the off-diagonal elements of this permuted correlation matrix, for each permutation, was used to generate a null-distribution to test the significance of the original correlations. P-values were calculated for each of the original correlations based on the frequency with which the elements of null-distribution were greater than or equal to the observed correlation coefficient. Note that the method of using the maximum of the permuted correlation matrix provides family-wise control for multiple comparisons (Westfall & Young 1993). Consequently, a nominal value of *P* < 0.05 was used as a cut off for statistical significance.

### The overlap between anatomical gyri and resting state functional clusters

For the overlap between the Desikan and Yeo atlases, the correspondence statistic was the normalized mutual information, NMI (Kvalseth 1987):

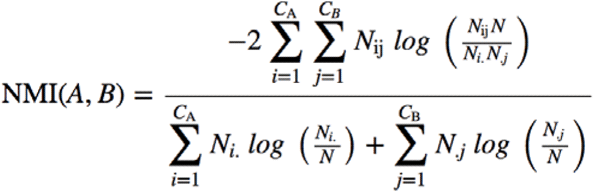

where A and B are the partitions, i.e., assignments of vertices to clusters (or regions) in the Desikan or Yeo atlas, respectively; C_A_ is the number of clusters (or regions) in partition A; C_B_ is the number of clusters (or regions) in partition B; N is the number of vertices, which is the same in both partitions; N_ij_ is the overlap between A’s cluster (or region) i and B’s cluster (or region) j, i.e. the number of vertices that they have in common; N_i_. is the total number of vertices in A’s cluster (or region) i; N._j_ is the total number of nodes in B’s cluster (or region) j; and this calculation follows the convention that 0×*log*(0)=0. The NMI ranges from 0 to 1, where 0 signifies that the partitions are totally independent and 1 that they are identical. See the discussion for an explanation of some of the terminology involved in the analysis of partitions, parcellations and functional clusters.

To test the statistical significance of the degree of the NMI between the Desikan atlas and the Yeo atlas, we generated 1000 rotational permutations of the data as described above. For each permutation, the NMI was re-estimated. The *P*-value was calculated as the frequency that which the permuted NMI estimates equaled or exceeded the actual NMI.

Additionally, we performed a *post hoc* analysis to determine which regions appeared to contribute disproportionately to the observed overlap. We generated a 34×7 confusion matrix, quantifying the overlap (in terms of total number vertices) between each region and each resting-state network. For each region and each network (41 total tests), the correspondence statistic was the pseudo-X^2^ testing whether each region was equally distributed between networks (weighted based on the total size of each network) and whether network was equally distributed between regions (weighted based on the size of each region). This statistic is called a pseudo- X^2^ to emphasize that it is not expected to be X^2^ distributed; rather, the null distribution is empirically determined by the permutation procedure. This analysis used FDR-correction for multiple comparisons correction (q<0.05).

### Overlap of meta-analytic patterns of activation with anatomical gyri and resting state functional clusters

For the test of overlap between the Yeo atlas and the 120 Neurosynth maps, the correspondence statistic was the pseudo-X^2^ transformation of the Wilk’s lambda from a Multivariate Analysis of Variance (MANOVA) (Krzanowski 1990). Significance of the observed correspondence statistic was determined with reference to the null distribution generated by 1000 rotational permutations of the data as described above.

As a *post hoc* analysis to investigate which of the 120 Neurosynth maps contributed disproportionately the global correspondence, we performed 120 separate tests where the correspondence statistics were the pseudo-F-statistics from Analysis of Variance (ANOVA) tests. Again, this statistic is called a pseudo-F to emphasize that it is not expected to have an F-distribution. The null distribution is determined by permutation. FDR correction was used to account for 120 comparisons.

The overlap between the Desikan atlas and the 120 Neurosynth maps was tested analogously, using MANOVA and post hoc ANOVA tests to generate correspondence statistics for spatial permutation tests.

## RESULTS

The permutation method allowed for rigorous hypothesis testing of the proposed relationships between patterns of functional activation, resting state fMRI networks and anatomical regions of interest. The probability density distribution appeared to stabilize by 1,000 permutations (Figure 1b-c), suggesting 1,000 as an appropriate number of permutations for subsequent statistical tests.

### Spatial Correspondences Between Metaanalytic Patterns of Activation Define Cognitive Clusters

The correlation coefficients between the 120 cognitive maps suggested a broad range in terms of the degree of relatedness between the maps (mean r = 0.004, sd = 0.12, range = −0.41 - 0.68). Hierarchical clustering of the maps revealed a non-trivial structure, for example with terms related to the language and reward systems, respectively, grouping together into relatively homogenous clusters of mutual correlation (Figure 2a). Of these correlations, 35 were statistically significant based on the family-wise correction for multiple comparisons (see Figure 2b). A network representation of these significant correlations (where nodes are cognitive terms and edges are significant correlations) revealed distinct components (edges within but not between sub-groups of nodes) corresponding to cognitive clusters including movement and motor planning, language, attention, memory, fear and reward (Figure 2b).

**Figure 2.**
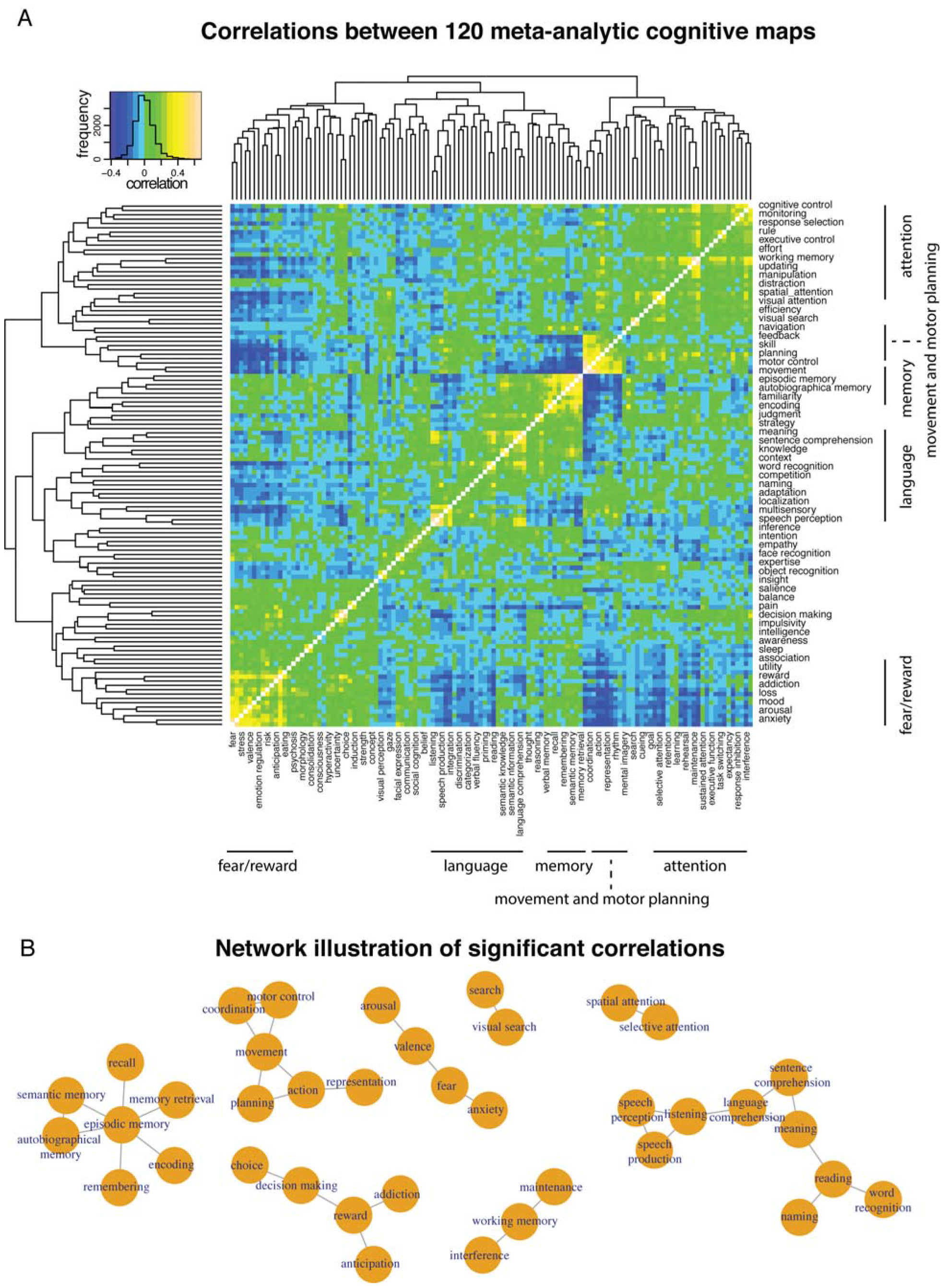
Correlation structure and significant correlation between meta-analytic activation patterns associated with 120 cognitive terms. A) Heat-map showing 120×120 correlation matrix. Terms or organized according to hierarchical clustering, with the resulting dendrogram shown to the top and to the left of the correlation matrix. Colors correspond to correlation coefficient, as shown in color key on top left. The color key also shows the frequency distribution of the correlations that comprise the matrix. Labels of the terms are shown to the right and to the bottom of the matrix, with the odd number labels shown on the bottom and the even number labels shown on the right (the order of the terms is “fear”, “anxiety”, “stress”, “arousal”, “valence,” etc.). B) Network illustration where the significant connections are illustrated as edges between the terms illustrated as nodes. The resulting network is comprised of 8 disconnected components; edges exist within each component’s nodes, but there are no edges between components.

### Gyral-Based Anatomical Regions of Interest Overlap with Resting-State Functional Networks

The hypothesized overlap between anatomical regions of interest and functional networks was supported by the permutation procedure. The actual NMI between the Desikan atlas (Figure 3a) and the Yeo atlas (Figure 3b), 0.389, is unlikely to be simply due to chance overlap (P=0.034) (Figure 3c). Put differently, patches of brain that form part of the same gyral-based region of interest were significantly more likely to also form part of the same intrinsic functional network. This overlap is not due simply to mutual spatial dependence, nor to mutual contralateral symmetry, as these potential confounds are controlled for by the permutation procedure. (The mean of the null distribution generated by the permutation procedure can be interpreted as an estimate of the contribution of these potential confounds to the observed NMI.)

*Post hoc* analysis suggested that this statistically significant correspondence resulted disproportionately from certain resting state networks and anatomical regions. Specifically, the visual, somatomotor and frontoparietal networks showed statistically significant correspondence with the gyral boundaries of anatomical regions after FDR correction for multiple comparisons (Table 1). Conversely, although no individual region’s statistical significance survived FDR correction for multiple comparisons, the anatomical regions that corresponded most closely with the boundaries of intrinsic resting-state networks were largely visual and motor areas, as evidenced by border-line significant *P*-values in Table 1.

**Table 1:**
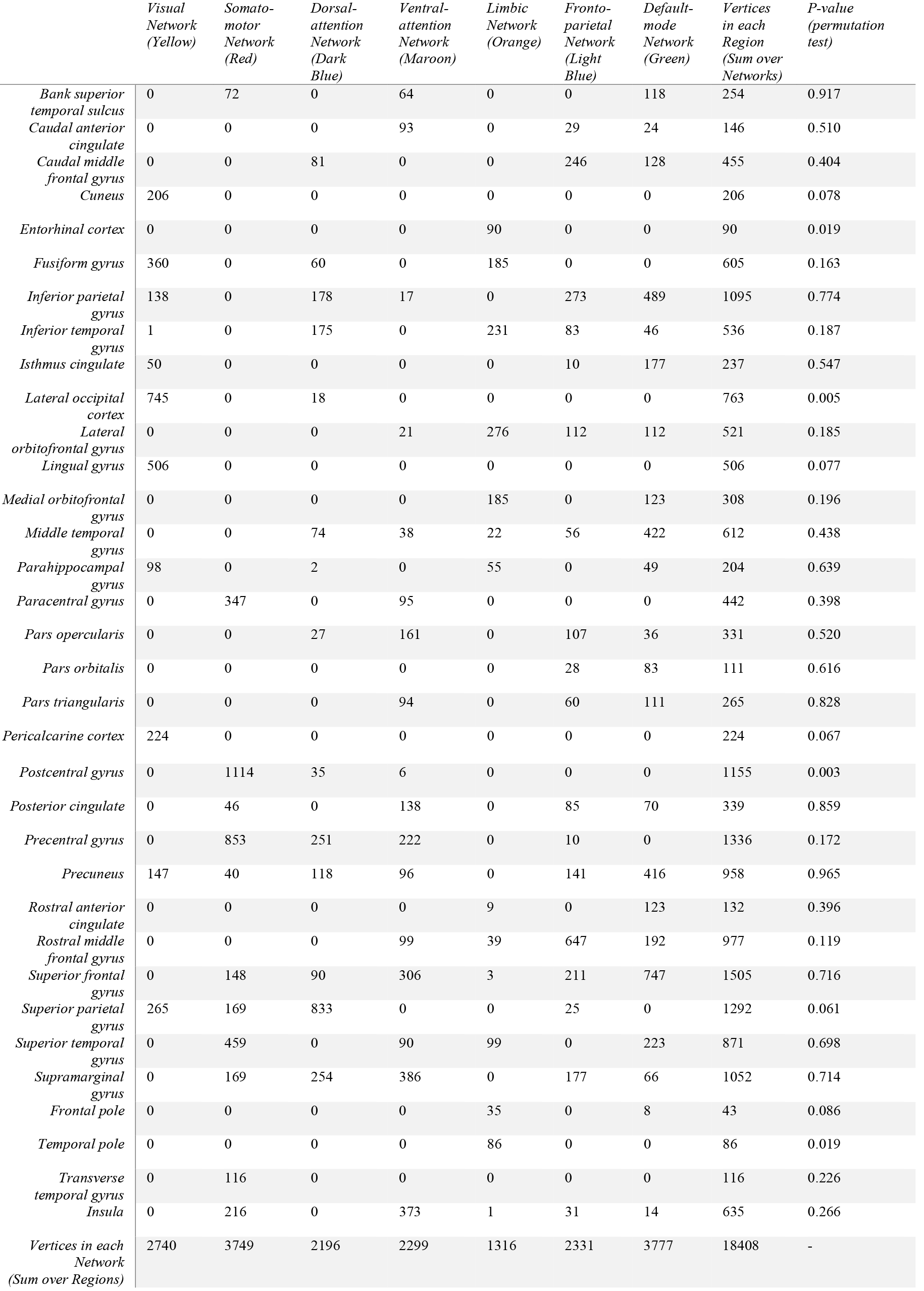
Confusion matrix between Desikan Atlas and Yeo Atlas, showing the number of vertices that are shared between each anatomical region and each resting state network. Labels of the networks are per the original description of the Yeo Atlas, while colors correspond to the Figure 3b. For networks, the correspondence statistic is the pseudo-X^2^ statistic for a test of whether each network is randomly distributed between reegions; while for regions, the correspondence statistic is the pseudo-X^2^ statistic for a test of whether each region is randomly distributed between networks. (These statistics are called pseudo-statics because they are not expected to be follow X^2^ distribution; rather, the *P*-values are determined based on the empirical distribution generated by the permutation procedure.) This *post hoc* test provides a sense of which regions and networks are contributing differentially to globally significant overlap between anatomical regions and functional networks (see Results). \**significant after FDR correction for multiple comparisons*

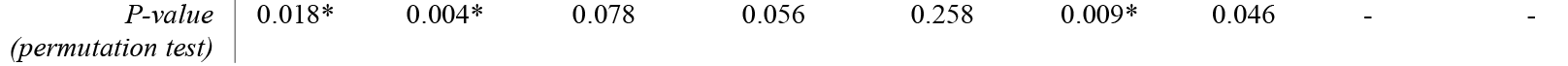

### Metaanalytic Patterns of Activation Overlap with Resting-State Functional Networks

Meta-analytic patterns of functional activation did not only cluster in space to define overarching cognitive domains, but also demonstrated statistically-significant overlap with functional networks defined by patterns of coordinated intrinsic cortical activity at rest (spatial permutation test using the pseudo-X^2^ transformation of the Wilk’s lambda from a MANOVA test as the correspondence statistic, *P*=0.001) (Figure 3e). Put differently, the patterns of activation shown by the meta-analyses of cognitive concepts tended to respect the boundaries of the resting-state functional networks. In the 120 *post hoc* permutation tests (using pseudo-F-statistics as the correspondence statistics), only “movement” was statistically significant after FDR-correction for multiple comparisons (*q*<0.05), but there was borderline significant correspondence with resting-state networks for the metaanalytic maps of working memory, pain, autobiographical memory and spatial attention (see illustrations in Figure 3d). Movement appeared to differentially activate the somato-motor network; working memory differentially activated the frontoparietal and dorsal attention networks; autobiographical memory differentially activated the default-mode network; and pain differentially activated the ventral attention network (see Table 1 for the labels of the individual resting-state networks).

### Metaanalytic Patterns of Activation Overlap with Gyral-Based Anatomical Regions of interest

Finally, as would be predicted based on their mutual overlap with intrinsic resting-state networks, metaanalytic patterns of activation also demonstrated statistically significant overlap with a classical parcellation of the cortical sheet defined by macroanamical gyral boundaries (spatial permutation test using the pseudo-X^2^ transformation of the Wilk’s lambda from a MANOVA test as the correspondence statistic, *P*=0.038) (Figure 3f). In other words, the patterns of activation shown by the metaanalyses of cognitive concepts tended to respect gyral boundaries. For the 120 *post hoc* permutation tests using pseudo-F-statistics as the correspondence statistics, only “listening” was statistically significant after FDR-correction for multiple comparisons (*q*<0.05), but there was borderline significant correspondence with gyral boundaries for the metaanalytic maps of movement, pain, speech perception and reward. Listening appeared to differentially activate superior temporal gyrus, tranverse temporal gyrus and the bank of the superior temporal sulcus; pain disproportionately activated insula, post central and caudal anterior cingulate cortex; movement differentially activated precentral, postcentral, supramarginal and superior parietal cortex; speech perception differentially activated superior temporal gyrus, bank of the superior temporal sulcus and transverse temporal gyrus; reward differentially activated the rostral anterior cingulate, lateral orbitofrontal and medial orbitofrontal gyrus.

**Figure 3.**
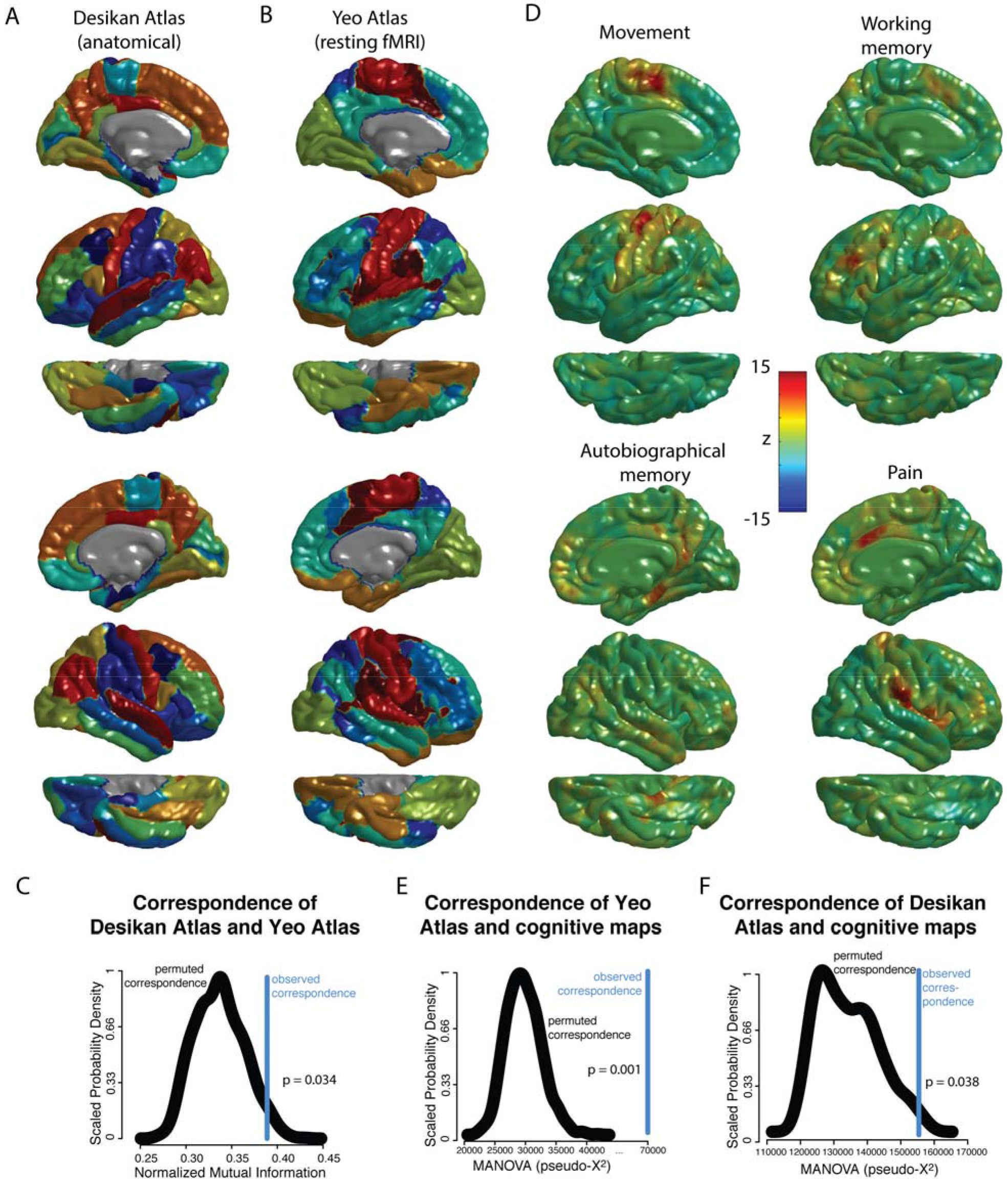
Spatial relationship between regions based on gyral landmarks (Desikan Atlas), intrinsic functional connectivity networks (Yeo Atlas), and task-based fMRI brain maps (Neurosynth meta-analyses). A) Representation of the Desikan Atlas, derived from manually identifying 34 in each hemisphere based on gyral landmarks, using 40 high resolutaion structural MRI scans. B) Representation of the Yeo atlas, derived by identifying 7 resting-state functional networks using a mixture model of 1000 resting-state fMRI scans. C) The normalized mutation information between the Yeo and Desikan Atlas, a measure of the similarity of the two atlases, for the original data as well as the probability density distribution of 1000 rotational permutations. The P-value is calculated as the frequency with which the permuted NMI equals or exceeds the actual NMI. D) Representation of 4 of the 120 brain maps derived from automated meta-analyses of cognitive concepts included in the cognitive atlas, with color scale corresponding to z-statistic (see methods). The top four cognitive terms are shown, ranked via F-statistic of 120 *post hoc* ANOVA tests of the relationship between these maps and the Yeo Atlas. As the maps are largely symmetric, for illustrative purposes, the left hemisphere is shown for movement and working memory, while the right hemisphere is shown for autobiographic memory and pain. E) The Chi-square transformation of the MANOVA test statistic where the networks of the Yeo atlas were the dependent variable and the 120 cognitive maps were the independent variables, for the original data as well as the probability density distribution of 1000 rotational permutations. The P-value was calculated as the frequency with which the permuted Chi-square statistic equaled or exceeded the actual test statistic. F) The Chi-square transformation of the MANOVA test statistic where the networks of the Desikan atlas were the dependent variable and the 120 cognitive maps were the independent variables, for the original data as well as the probability density distribution of 1000 rotational permutations. The P-value was calculated as the frequency with which the permuted Chi-square statistic equaled or exceeded the actual test statistic.

## DISCUSSION

Our work addresses a number of methodological issues that arise in the statistical comparison of brain maps, and in doing so provides evidence in support of specific biological hypotheses regarding functional topography and function-structure relationships in the human cerebral cortex. We advocate a spatial permutation approach to the issue of comparing cortical maps. This approach demonstrates partial overlap or convergence between meta-analytic maps of functional activations, gyral-based anatomical regions of interest, and resting-state functional connectivity networks. In making the code for these analyses available and discussing the methodological issues, we hope to draw attention to this issue and increase the statistical rigor with which “the correspondence problem” is approached.

The utility of the rotational permutation approach is illustrated by the inter-relationships between the anatomical substrates of 120 cognitive functions. Although the 120×120 correlation matrix appears to have meaningful structure (Figure 2a), P-values generated from a parametric test of the significance of these correlation coefficients would have extremely high false positive rates, because of falsely assuming spatial and contralateral independence. (Non-parametric tests and permutation procedures that do not preserve spatial and contralateral dependence would have similarly high false positive rates.) In addition, a correction for multiple comparisons across the 7140 correlations is required, but as these correlations are not independent, standard approaches for multiple comparisons correction may be too conservative. The permutation procedure addresses both of these concerns, by generating permuted data with the same spatial and contralateral structure of the original data, as well as a null distribution of maximum correlation coefficients that allows for a straightforward family-wise correction for multiple comparisons. In general, there is an analogy between spatial permutation and other well-described (although perhaps under-utilized) permutation approaches (E. T. Bullmore et al. 1999; Nichols & Holmes 2002; Winkler et al. 2014); the spatial coordinates of the maps are permuted, rather than the task/rest labels of a functional scan, or the patient/control labels in a case-control study.

Spatial dependence and contralateral symmetry are multi-faceted and sometimes problematic issues in neuroimaging studies. On the one hand, there is ample evidence that spatial constraints are a biological principle of brain network organization (E. Bullmore & Sporns 2012). On the other hand, spatial dependence is also introduced by image processing pipelines, for example, when images are smoothed in order to make statistical comparisons between anatomically divergent individuals. In addition, motion artifact may introduce artefactual smoothing into both functional (Satterthwaite et al. 2012) and anatomical (Alexander-Bloch et al. 2016) scans. Regardless of the underlying source of spatial dependence, or whether it is biological or artefactual in a given experimental context, it has the potential to confound statistical inference about partial overlap or convergence between brain maps. Contralateral symmetry is more generally appreciated as “biological” compared to spatial autocorrelation within hemispheres. However, confounds such as motion are likely to similarly impact contralateral homologues (which are the same distance from the axes of rotation). Therefore, similarly to spatial dependence, symmetry can falsely inflate the apparent overlap when comparing brain maps that include both left and right hemisphere data.

There is no single accepted methodology to compare brain parcellations and partitions, e.g., maps of anatomical regions of interest and or intrinsic functional connectivity networks. The terminology can be confusing: it is common to call a map of anatomical regions a parcellation (Van Essen et al. 2012), while maps of intrinsic connectivity (or other kinds of connectivity) are described as partitions, communities, clusters or modules (Alexander-Bloch et al. 2010). However, the practical distinction between these two kinds of maps is that anatomical regions, by definition, form spatially contiguous patches. Partitions, while often spatially contiguous, *may* be spatially distributed; in functional parcellations such as the Yeo atlas, sensorimotor clusters tend to form distinct patches, while spatially discontiguous clusters are distributed across association cortical areas. For the purposes of statistical comparison, similar statistics can quantify the degree of overlap, such as the adjusted Rand index (Hubert & Arabie 1985), a variation of information criterion (Meilă 2007), and normalized mutual information (Kvalseth 1987). Because properties such as the number of clusters and the degree of spatial dependence affect these statistics, they cannot be interpreted simply based on their magnitude. Spatial permutation provides a framework for the significance of these statistics to be evaluated.

By leveraging these strengths of spatial-permutation testing for the comparison of three distinct types of cortical map, our study provides a number of insights into cortical organization. First, we provide quantitative evidence that task-induced patterns of brain activation (Neurosynth meta-analytic maps) are non-randomly related to patterns of coordinated brain activity at rest (Yeo 7-network resting state parcellation). The ability to test the apparent similarity between the two modes of coordinated brain activity (Eickhoff et al. 2017) provides a necessary empirical and technical foundation for asking how this similarity arises, and whether inter-individual differences in the strength of this spatial coupling are relevant for behavioral differences in health and disease (Braga & Buckner 2017). Second, we test the spatial inter-dependence between functional and macroanatomical topography of the cortical sheet. The non-random organization of brain function with respect to gyral features is evident with respect to both coordinated brain activity at rest (Yeo Atlas) and task-evoked brain activity (Neurosynth data), and varies in strength across different functional systems. Specifically, we find that the degree of alignment between function and structure tends to be stronger for primary input/output systems (visual, auditory, motor) than for higher-order associative systems. The corresponding gyral features that show closest alignment for the spatial organization of brain activity during rest and tasks are aligned with those “primary sulci” that arise earliest in prenatal human brain development (Nishikuni & Ribas 2013), show least variance in morphology across individuals (Mangin et al. 2004) and most consistent correspondence with the boundaries of cytoarchitectonically-defined cortical areas (Fischl et al. 2008). These observations suggest the operation of a strong conjoint constraint on the anatomical and functionally patterning of lower-order cortical regions, such that these display an alignment of gyral and functional boundaries that can be detected at the group level. The spatial permutations approach represents a promising quantitative framework for moving beyond group-level analyses in future work to probe structure-function relationships in individual cortical sheets.

Several methodological issues with the present analysis should be noted. First, although the present method is limited to cortical surface maps, it could be extended to sub-cortical or volumetric maps, most readily for structures such as the thalamus that could reasonably be projected onto a spherical volume. Second, to highlight the methodological approach, we have chosen to present biological results that are largely confirmatory in nature. Clusters of similar functional maps correspond to *a priori* similar cognitive functions, which plausibly explains the correspondence of the meta-analytic maps. The inter-dependence of resting state networks and multivariate patterns of functional activation has previously been described using different methodological approaches (Biswal et al. 1995; Smith et al. 2009). Finally, some degree of overlap between known anatomical regions of interest and functional connectivity networks is generally assumed, but this assumption has not previously been formally tested to our knowledge. The illustrative analyses also depend on upstream methodological choices. For example, the choice of terms to include in the automated meta-analysis clearly constrains downstream results, and different methods of controlling for multiple comparisons could result in different results.

Despite these limitations, the spatial permutation methods described, applied and disseminated by our study constitute a useful advance upon current methods for the comparison of cortical maps, and thereby open up new possibilities for the surface based integration of diverse neuroimaging data.

